# A comparative study of neuroendocrine heterogeneity in SCLC and NBL

**DOI:** 10.1101/2022.11.17.516959

**Authors:** Ling Cai, Ralph J. DeBerardinis, Yang Xie, John D. Minna, Guanghua Xiao

**Affiliations:** Quantitative Biomedical Research Center, School of Public Health, UT Southwestern Medical Center, Dallas, TX 75390, USA; Children’s Research Institute, UT Southwestern Medical Center, Dallas, TX 75390, USA; Simmons Comprehensive Cancer Center, UT Southwestern Medical Center, Dallas, TX 75390, USA; Howard Hughes Medical Institute, University of Texas Southwestern Medical Center, Dallas, TX 75390, USA; Department of Bioinformatics, University of Texas Southwestern Medical Center, Dallas, TX 75390, USA; Hamon Center for Therapeutic Oncology Research, University of Texas Southwestern Medical Center, Dallas, TX 75390, USA; Department of Pharmacology, UT Southwestern Medical Center, Dallas, TX 75390, USA; Department of Internal Medicine, University of Texas Southwestern Medical Center, Dallas, TX 75390, USA

## Abstract

Lineage plasticity has long been documented in both small cell lung cancer (SCLC) and neuroblastoma (NBL), two clinically distinct neuroendocrine (NE) cancers. In this study, we quantified the NE features of cancer as NE scores and performed a systematic comparison of SCLC and NBL. We found NBL and SCLC cell lines have highly similar molecular profiles and shared therapeutic sensitivity. In addition, NE heterogeneity was observed at both the inter- and intra-cell line levels. Surprisingly, we did not find a significant association between NE scores and overall survival in SCLC or NBL. We described many shared and unique NE score-associated features between SCLC and NBL, including dysregulation of Myc oncogenes, alterations in protein expression, metabolism, drug resistance, and selective gene dependencies. Our work establishes a reference for molecular changes and vulnerabilities associated with NE to non-NE transdifferentiation through mutual validation of SCLC and NBL samples.

## Introduction

Small cell lung cancer (SCLC) and neuroblastoma (NBL) are two very different cancer types with respect to their etiology, mutation spectrum/load, classification scheme, therapeutic strategy, and prognosis. SCLC, accounting for 13% of lung cancers, is predominantly found in heavy smokers, with almost ubiquitous co-mutation of *RB1* and *TP53*, and has a five-year survival rate of 7% as the disease is highly metastatic and two-thirds of the patients are diagnosed at the extensive stage. SCLC bears much resemblance to pulmonary neuroendocrine cells in their morphology and expression of NE markers (Bensch et al., 1968), but studies from genetically engineered mouse models suggest that some SCLC may also arise from other lung cell types (Yang et al., 2018). NBL, accounting for 6% of childhood cancers in the US, is derived from sympathoadrenal progenitor cells within the neural crest (Johnsen et al., 2019), often develops in and around the adrenal gland, exhibits frequent genetic alterations in *MYCN* or *ALK*, and has a five-year survival rate of 81%. Despite these differences, both SCLC and NBL are neuroendocrine (NE) tumors, and NE markers are routinely used in immunohistochemistry (IHC) to facilitate the clinical diagnosis of both cancer types. As one of the “small round blue cell tumors” of childhood, undifferentiated NBL also highly resembles SCLC histologically.

Interestingly, the ability to transdifferentiate from the NE to non-NE lineage has been documented for both SCLC and NBL. Over 35 years ago, “classic” (NE) and “variant” (non-NE) SCLC were reported based on distinct cellular morphologies and biochemical properties (Gazdar et al., 1985). In the recent decade, studies have shown that transdifferentiation of SCLC gives rise to intratumoral heterogeneity and mediates chemoresistance (Calbo et al., 2011; Lim et al., 2017). More recently, it was shown that REST, YAP, and NOTCH mediate NE transition in both SCLC and normal lung (Shue et al., 2022). For NBL, morphologically distinct cell types from cell lines established from the same patient tumor were observed over 50 years ago (Tumilowicz et al., 1970). Distinct biochemical properties and the ability to interconvert have been reported for isogenic cell subclones (Ross et al., 1983). In two more recent studies, the “sympathetic noradrenergic”(NE) and “neural crest cell-like” (non-NE) (Boeva et al., 2017), or “adrenergic” (NE) and “mesenchymal” (non-NE) (van Groningen et al., 2017) NBL cell states have been shown to exhibit distinct epigenetic and transcriptomic profiles. It has also been shown that NOTCH regulates TF networks to drive NE transition in NBL and contribute to the development of chemoresistance in NBL (van Groningen et al., 2019). These independent studies converged on similar NOTCH-mediated mechanisms in NE lineage switch and suggest shared NE-associated properties across different cancer types. However, the extent of such similarity is still unclear. In this study, we re-analyzed the molecular and clinical data generated from SCLC and NBL cell lines and tumors to compare their associations with NE heterogeneity side-by-side, to reveal the concordance and idiosyncrasy in the landscape of NE state-associated features in both cancer types.

## Results

### NBL and SCLC cell lines are molecularly similar

We have previously established an NE score calculation method for SCLC samples based on a gene expression signature generated from SCLC cell line transcriptomic data (Cai et al., 2021a; Zhang et al., 2018). This method takes the expression data of 25 NE genes and 25 non-NE genes as inputs and assigns a score ranging from −1 (non-NE) to 1 (NE) to each sample. From the pan-cancer study cancer cell line encyclopedia (CCLE)/dependency map (DepMap) (Tsherniak et al., 2017) RNA-seq dataset, we averaged the expression of these 50 genes by different cancer lineages and performed hierarchical clustering (**Figure 1a**). Among the NE cancer types in the sub-cluster with high expression of NE genes, SCLC and NBL had the highest number of cell lines in the CCLE collection. This allowed us to leverage the multi-dimensional profiling data from CCLE and DepMap for an in-depth comparison between SCLC and NBL. We computed NE scores for the pan-cancer cell lines and clustered the cell lines based on transcriptomic, functional proteomic (based on Reverse Phase Protein Arrays, RPPA), metabolomic, gene dependency, and drug sensitivity features (**Figures 1b-h**). We observed tight clusters of high-NE-score cell lines and tight clusters of SCLC and NBL cell lines in each clustering analysis. These results suggest that SCLC and NBL cell lines are highly similar in these molecular aspects.

**Figure 1.**
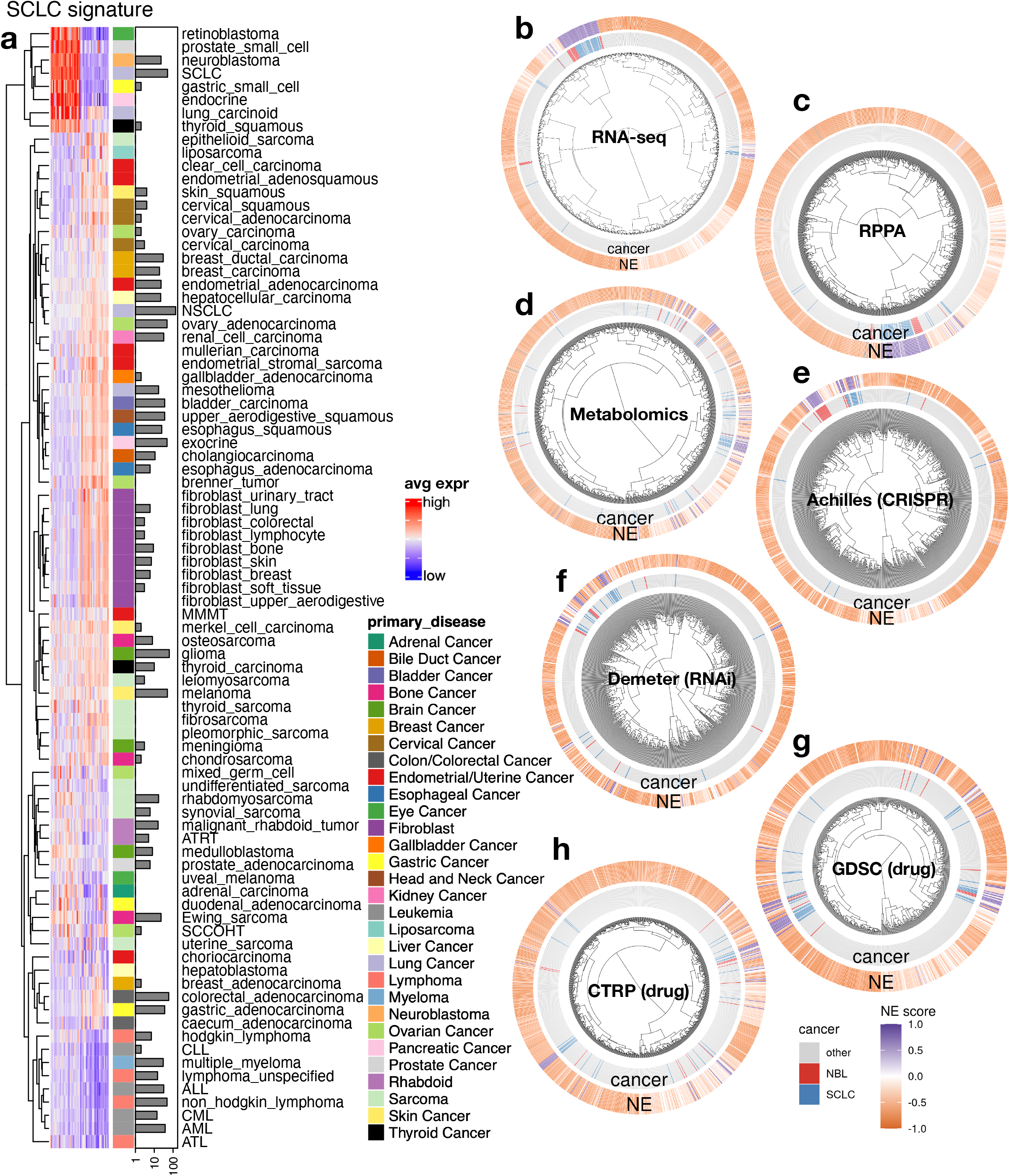
The molecular similarity between NBL and SCLC. **a**. SCLC NE signature gene expression across different cancer lineages. The expression of NE (left half) and non-NE (right half) genes were averaged by cancer lineages and plotted as a heatmap. The per-lineage cell line counts are visualized as bars plotted right to the heatmap. Note that SCLC and NBL are the two cancer types with the highest number of cell lines in the cluster with high expression of NE genes. **b**-**h**, hierarchical clustering of 1165 cancer cell lines by expression of 19159 genes (**b**), 897 lines by 214 RPPA features (**c**), 926 lines by 225 metabolites (**d**), 688 lines by CRISPR effect score of 509 genes (**e**), 648 lines by RNAi effect score of 375 genes (**f**), 624 lines by 208 compounds from GDSC (**g**), and 794 lines by 168 compounds from CTRP (**h**). Note the clustering for RNA-seq data was based on the top 10 principal components, RPPA and metabolomics clusterings were based on all available features; dependency and drug clusterings were based on selected consistent features as previously summarized.

### NE heterogeneity can be observed at inter- and intra-cell line levels for both SCLC and NBL

We ranked cell lines in the CCLE panel based on their NE scores and observed that most of the SCLC and NBL cell lines were enriched in the top, with positive NE scores. However, we also found a few SCLC and NBL lines with negative NE scores, revealing inter-cell line NE heterogeneity, as previously reported (**Figure 2a**).

**Figure 2.**
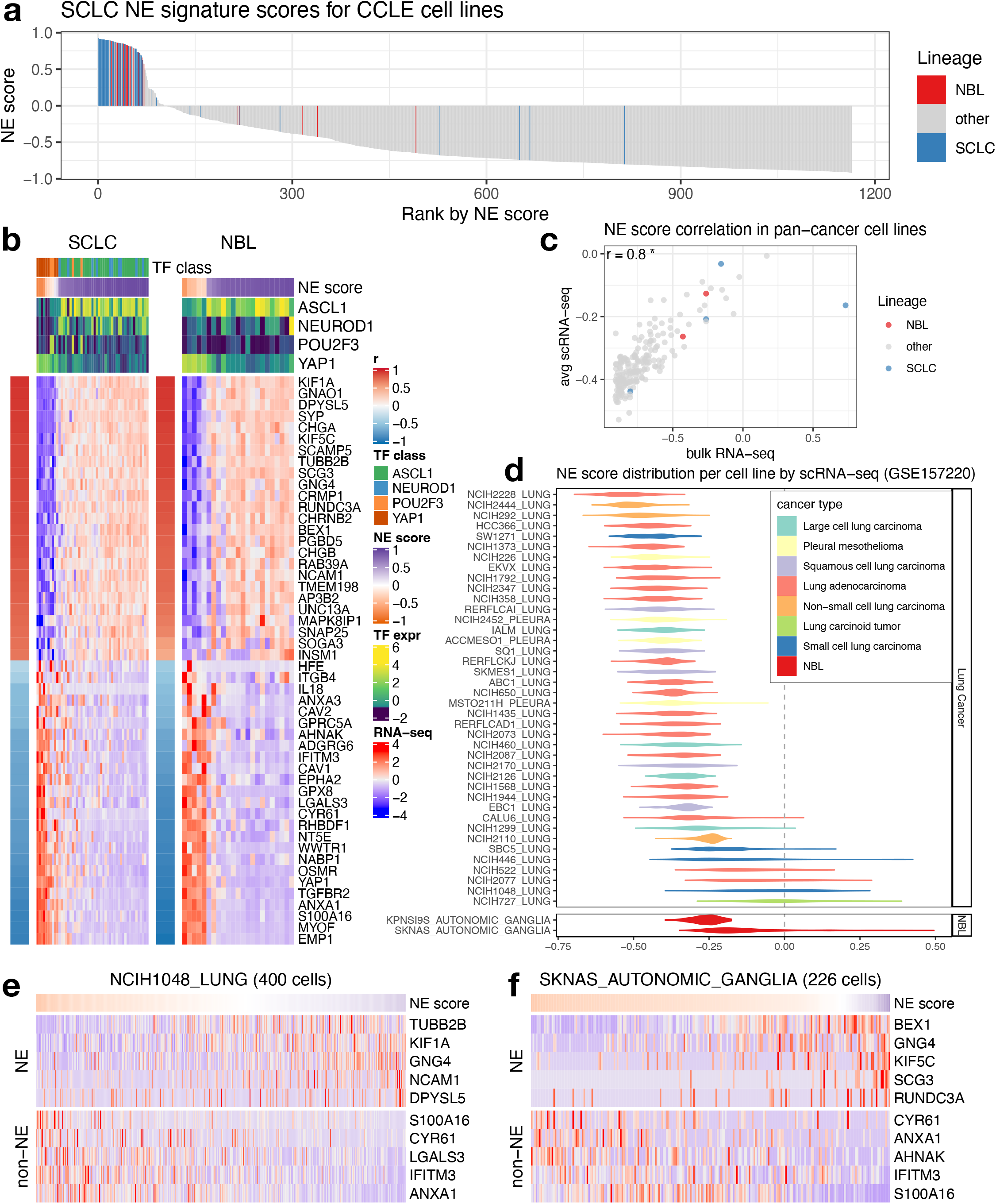
Inter- and intra-cell line NE heterogeneity. **a**. Inter-cell line NE heterogeneity. NE scores for CCLE pan-cancer cell lines were ranked from high to low. SCLC and NBL lines were highlighted by colors. Although most of the SCLC and NBL lines have high NE scores, a few of them also have low NE scores. **b**. Consistent gene expression pattern for SCLC NE signature genes observed for SCLC and NBL cell lines. Cell lines are in columns. Red/blue column left to the heatmap annotates the correlation between the gene expression and NE score; the expression of SCLC driver transcription factors (TFs) and NE scores was annotated above the heatmap. SCLC lines were further classified into four transcription factor (TF) classes. **c**. Average NE scores from scRNA-seq data align well with NE scores from bulk RNA-seq data for pan-cancer cell lines. **d**. Distribution of NE scores for lung cancer and neuroblastoma cell lines based scRNA-seq data. **e**-**f**. Intra-cell line NE heterogeneity. High- and low-NE score cells are found to coexist within the same SCLC cell line NCI-H1048 (**e**) or NBL cell line SKNAS (**f**). Single cells are in columns. Due to the high dropout rate of scRNA-seq data, only the top abundantly expressed genes are visualized.

We examined the expression of SCLC NE score signature genes in SCLC and NBL cell lines. Although the signature was established in SCLC cell lines, it was also highly differentially expressed in NBL cell lines (**Figure 2b**). Four key transcription factors (*ASCL1, NEUROD1, POU2F3*, and *YAP1*) have been proposed to define the four molecular subtypes of SCLC. We examined their expression in NBL cell lines and observed similar segregation by NE score. While high-NE-score NBL lines were found to have high expression of *ASCL1* or *NEUROD1*, low-NE-score NBL lines had high expression of *YAP1*. However, no NBL line was found to express high levels of *POU2F3*, a tuft cell regulator (Huang et al., 2018) (**Figure 2b**). These results suggest that similar transcriptional regulations are involved in driving NE heterogeneity in SCLC and NBL cell lines. Using scRNA-seq data available for a panel of pan-cancer cell lines (Kinker et al., 2020), we compared the scRNA-seq-based average NE score to the bulk RNA-seq-based NE scores for 191 cell lines and found a strong correlation (**Figure 2c**). Interestingly, when we examined the distribution of NE scores for lung cancer and NBL cell lines, we observed that some SCLC and NBL cell lines had broader NE score distributions than others (**Figure 2d**). Using scRNA-seq data from SCLC patient tumors, we could observe the coexistence of high- and low-NE-score SCLC cells within the tumors that exhibited highly variable NE scores (**Figure S1**). Upon close examination of NE and non-NE gene expression across single cells in the SCLC cell line NCI-H1048 and NBL cell line SKNAS, we also observed the coexistence of high- and low-NE score cells within the same cell line (**Figures 2e-f**). These results suggest that the lineage heterogeneity observed in patient tumors is preserved both among cell lines and within individual cells from the same cell line.

### NE scores do not associate with overall survival in SCLC or NBL

Next, we tested whether NE scores were associated with disease outcomes in SCLC and NBL. As most patients with SCLC are diagnosed at an extensive stage, surgical resection of SCLC primary tumors is rare in practice. A recent study that profiled biopsied metastatic SCLC samples found no association between NE score and outcome (Lissa et al., 2022). We also investigated the prognosis association in a previously published dataset generated from 81 surgically resected SCLC tumors, of which 30 are stage III-IV samples (George et al., 2015). We also did not find a significant association between the NE scores and overall patient survival (**Figure 3a**). With multiple NBL tumor datasets available, we performed a meta-analysis to assess the association between NE scores and overall survival in NBL. We also did not observe a consistent and significant result (**Figure 3b**). In the NBL datasets we investigated, the previously reported prognostic factors – age, MYCN amplification, and INSS stage 4 disease–were consistently associated with worse overall survival (**Figure S2a-c**), but we did not observe a significant difference in NE scores in groups stratified by these factors (**Figures S2d-f**). A small effect size was observed for NE score difference by relapse/progression status (**Figure S2g**); however, when comparing paired naïve and relapse samples from the same patient in two independent NBL studies, we did not identify a statistically significant difference in NE scores (**Figure 3c**). These findings suggest NE scores are not associated with prognosis in SCLC or NBL.

**Figure 3.**
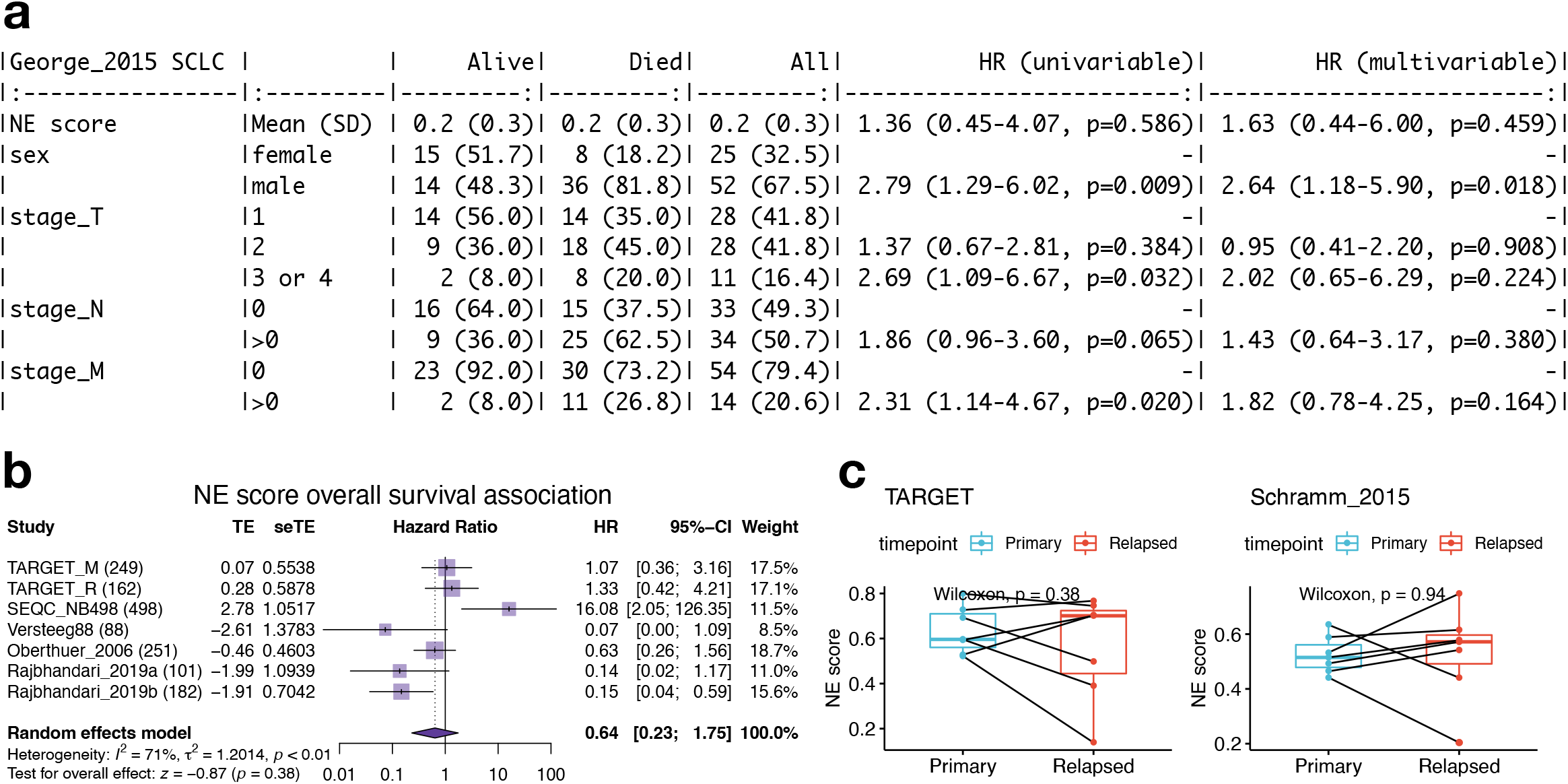
NE score is not associated with overall survival in SCLC or NBL. **a**. Survival association analysis for SCLC based on 79 patients from the George_2015 study. NE score is not significantly associated with overall survival in univariate Cox regression or a multivariate model controlling for Sex and TNM stage. **b**. Meta-analysis for NBL based on seven studies and 1,531 patients. The result is also not statistically significant although significant results could be observed for individual studies, the trend was different.**c**. NE scores are not significantly altered in NBL relapsed samples. Paired samples from the same patients in two independent studies were compared.

### Myc oncogenes are differentially activated by NE states in SCLC and NBL

As members of the Myc oncogene family (*MYC*, *MYCN*, and *MYCL*) have been implicated in SCLC and NBL oncogenesis (Huang and Weiss, 2013; Johnson et al., 1987), we attempted to dissect their relationship with the NE state. First, we examined copy number alterations of Myc oncogenes (**Figure 4a**). We found that *MYC* and *MYCL* were enriched in high-NE-score SCLC lines, whereas *MYCN* amplification was enriched in high-NE-score NBL lines. As MYCL is located on chromosome 1p, a frequently deleted region in NBL, *MYCL* loss appears to be frequent in NBL lines (**Figure 4a**). Examination of the gene expression data showed that the patterns for *MYCL* and *MYCN* agreed well with the copy number data (**Figure 4b**). Having made these observations in cell lines, we further examined the transcriptomic data from multiple SCLC and NBL studies. For SCLC, we included our in-house cell line RNA-seq data (UTSW cell lines), PDX dataset (Drapkin_2018), and four tumor datasets (**Figures 4c**). For NBL, we included three more cell line datasets along with the CCLE RNA-seq data (**Figures 4d**) and assembled 11 tumor datasets (**Figures 4e**). Meta-analyses with these datasets verified that the NE score associations with Myc oncogenes were consistent between multiple cell lines (**Figure 4b**) and patient tumor datasets (**Figure 4f**). Combined analysis of copy number, gene expression, and NE scores in the CCLE cell line dataset revealed upregulation of MYC expression in the low-NE-score lines without copy number gain, suggesting the transcriptional activation of *MYC* expression in the non-NE state for both SCLC and NBL (**Figure 4g**). We retrieved MYCN amplification status from eight NBL tumor datasets and assessed the association between NE score and MYCN expression while controlling for MYCN amplification (**Figure 4h**). Much stronger associations were observed across multiple studies in this multivariate linear model (**Figure S3**), suggesting the transcriptional activation of *MYCN* expression in the NE state NBL tumors. In summary, SCLC and NBL exhibit not only differential copy number gains but also differential transcriptional regulation for MYC family genes with regard to their NE status, with MYC transcriptionally upregulated in the non-NE state, MYCL preferentially amplified in high-NE score SCLC, and MYCN preferentially amplified and transcriptionally upregulated in high-NE score NBL.

**Figure 4.**
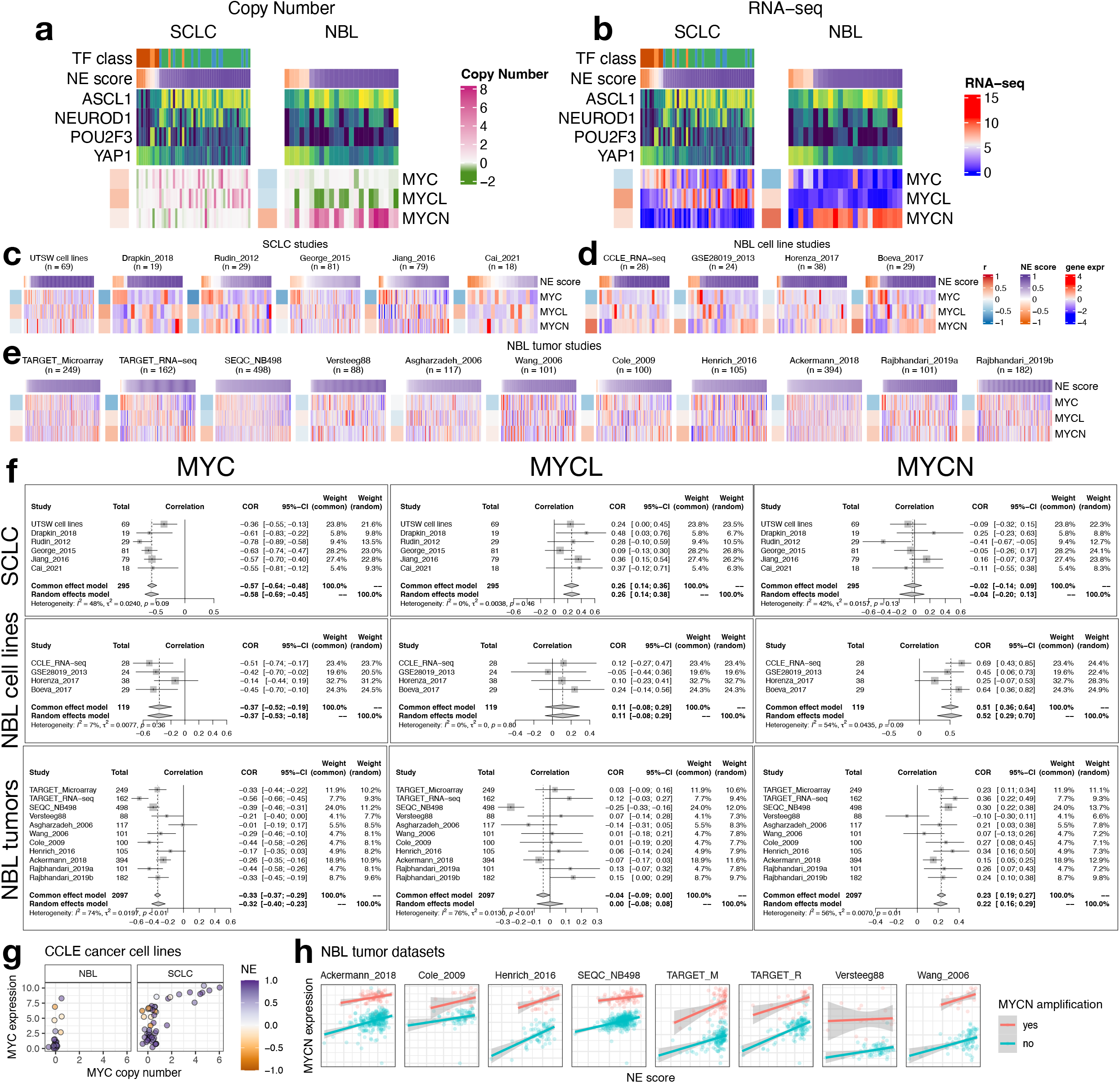
NE score association with members of the Myc oncogene family in SCLC and NBL. **a**-**b**. Copy number (**a**) and RNA expression (**b**) of Myc family genes in SCLC and NBL cell lines. Note that although MYC amplification was higher in the high-NE-score SCLC cell lines, its gene expression was higher in the low-NE-score cell lines for both SCLC and NBL lines. Frequent *MYCL* loss was found in NBL because *MYCL* is located in a frequently deleted region (chromosome 1p) in NBL. **c**. NE score vs. Myc gene member expression in SCLC studies. “UTSW cell line” is a cell line dataset; “Drapkin_2018” is a patient-derived xenograft dataset; “Rudin_2012”, “George_2015”, “Jiang_2016” and “Cai_2021” are all patient tumor datasets. **d**-**e**. NE score vs. Myc gene member expression in NBL cell line datasets (**d**) and tumor datasets (**e**). Note that some of the same cell lines were profiled in multiple studies. **f**. Forest plots visualizing meta-analysis of NE score association with Myc family genes. *MYC* expression is consistently associated with lower NE scores in SCLC and NBL samples (left). *MYCL* expression positively correlates with NE scores in SCLC but not NBL samples (middle). *MYCN* expression positively correlates with NE scores in NBL but not SCLC samples.

### Consistent proteomic and metabolic changes are associated with NE-to-non-NE transition in SCLC and NBL

We performed NE score correlations with 12 sets of data from the CCLE/DepMap studies (**Tables S1-S12**). These include four sets of omics data (miRNA, histone PTM, RPPA, and metabolomics), six sets of compound screening data (CCLE, CTRP, GDSC1, GDSC2, PRISM_1ST, and PRISM_2nd), and two sets of gene dependency screening data (Demeter for RNAi and Achilles for CRISPR). The overall NE score association concordance was quite good for the omics datasets (**Figure S4**).

Upon close examination of the results of the RPPA associations (**Figure 5a**), we found most of the NE-score-associated features identified in SCLC cell lines could also be observed in NBL cell lines. Among the exceptions, Rb protein is decreased in the high-NE score SCLC, leading to an increase in cyclin E2 but this was not observed in NBL lines (**Figure 5c**), which could be explained by the frequent *RB1* loss that occurs in SCLC but not NBL. Although a previous study suggests *RB1* loss is highly enriched in YAP^off^ small-cell/neuro/neuroendocrine cancer lineages (Pearson et al., 2021), the absence of *RB1* mutation in NBL suggests the existence of an Rb-independent mechanism for YAP inactivation in NBL. Interestingly, in both SCLC and NBL, CDK-interacting protein/kinase inhibitory protein (CIP/KIP) p21 is upregulated in the low-NE-score lines whereas another CIP/KIP p27 is downregulated, suggesting that the NE state-specific cell cycle regulators are still consistent in these two cancer types despite difference in the upstream Rb loss. In the low-NE-score lines of both SCLC and NBL, we observed higher levels of receptor tyrosine kinases and their phosphorylation (EGFR, EGFR_pY1068, HER2_pY1248, and VEGFR2), higher levels of Hippo signaling components (YAP, YAP_pS127, and TAZ), pro-inflammatory proteins (p62, NF-kB-p65_pS536, PAI-1, and annexin 1), ribosome biogenesis markers (S6_pS240_S244 and S6_pS235_S236), and cell adhesion proteins (paxillin and CD49b). In the high-NE score lines of both SCLC and NBL, we found higher apoptotic machinery components (Smac, Bcl-2, Bim, and Bax), DNA repair proteins (MSH2 and MSH6), translation inhibitor 4E-BP1, and microtubule regulator Stathmin. Unique to SCLC, we observed higher epithelial junction proteins (Claudin-7 and E-cadherin) in the high-NE-score lines, suggesting that NE SCLC lines are of the epithelial lineage, whereas NE NBL lines are not (**Figure 5a**). We also examined the metabolomic associations and observed similar consistency between SCLC and NBL lines. (**Figure 5b**) In particular, many cholesteryl esters (CEs) were found to have higher levels in the low-NE-score SCLC lines; a weaker but similar trend was observed in NBL lines. We also found that both SCLC and NBL low-NE-score cell lines exhibited higher levels of citrate, aconitate, and isocitrate, three interconvertible metabolites, through the action of aconitase (**Figures 5b** and **d**).

**Figure 5.**
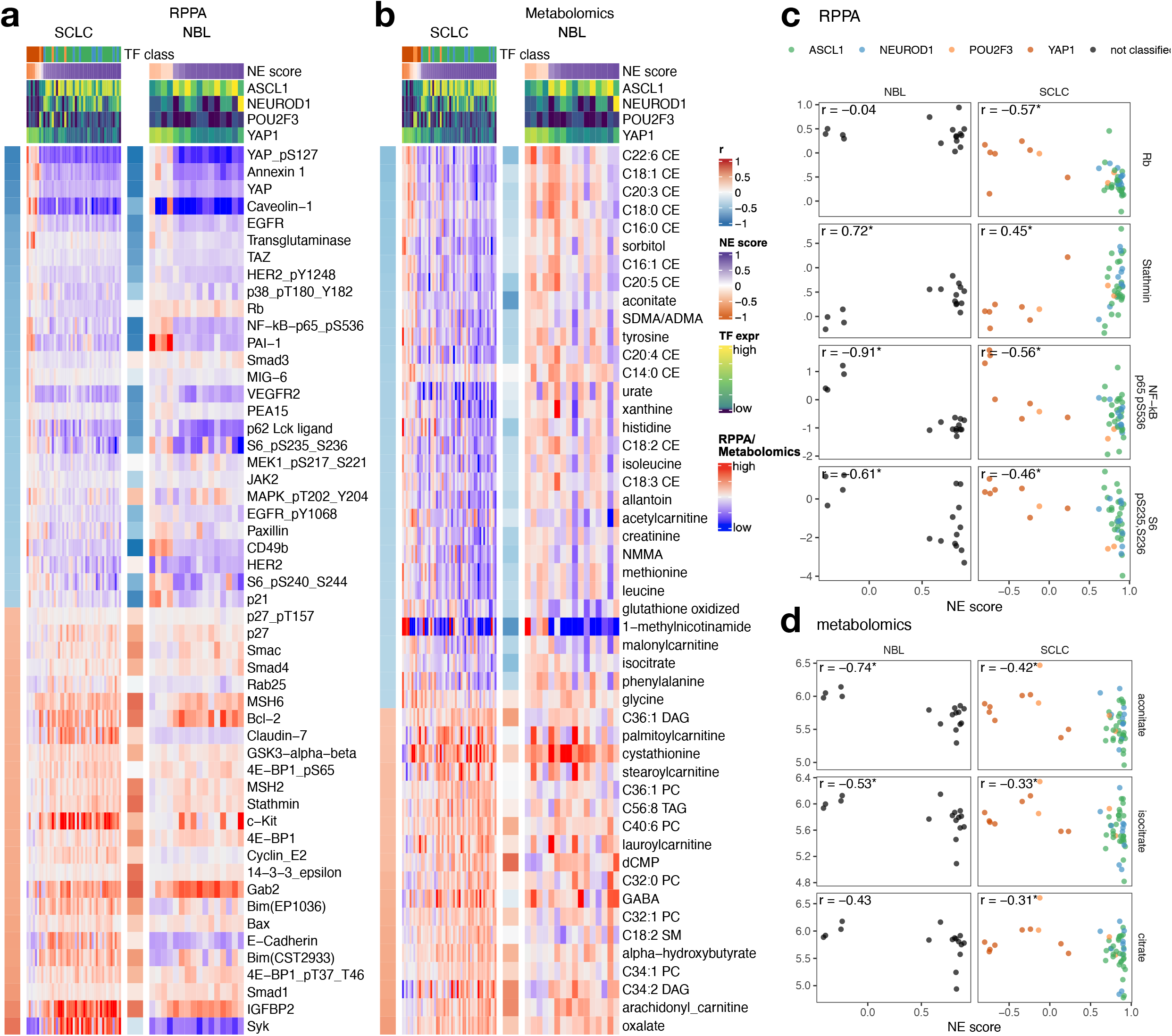
NE score-associated protein and metabolic features are largely consistent in SCLC and NBL cell lines. **a-b**. Heatmaps visualizing the relationship between NE scores and selected functional proteomic feature (**a**) or metabolites (**b**). In each heatmap, the left-side column denotes the Pearson correlation between the selected feature on the row and the NE score. The top colored rows denote NE scores and SCLC TF expression. The features were selected based on NE score correlation from the SCLC cell lines, adjusted p-value (p.adj) < 0.05 was used to select RPPA features and p.adj < 0.1 was used to select metabolic features. Note that although the selection was made from SCLC cell lines, a very similar pattern could be observed in NBL cell lines. **c**-**d**. Scatterplots visualizing the relationship between selected RPPA (**c**) and metabolic (**d**) features and NE scores in NBL and SCLC cell lines.

### Consistent and unique therapeutic vulnerabilities in NE and non-NE subtypes of SCLC and NBL

The SCLC vs. NBL concordance for NE score – drug sensitivity associations was poorer than the omics data (**Figure S4**). We have previously demonstrated that drug screening data are more consistent for compounds directed against functionally important targets that are differentially expressed in a panel of cell lines (Cai et al., 2021b). For many of the compounds included in the screens, their targets may not be functionally important in the small panel of cell lines tested, which may explain the overall lower consistency. We reviewed the results (**Tables S5-S10**) to identify the most consistent associations across the multiple compound screens. Nine classes of compounds with different mechanisms of action (MOA) were selected. For each MOA class, we compared the NE score associations for different compounds in NBL and SCLC (**Figure 6a**). We also used meta-analysis to generate a summary correlation coefficient for each class of compounds from the SCLC and NBL assessments (**Figures S5** and **6b**). We found that in both SCLC and NBL, cell lines with higher NE scores were more resistant to drugs that target MEK, mTOR, XIAP, LCK, HSP90, and Abl but were more sensitive to BCL inhibitors. We also observed that higher NE scores were associated with resistance to microtubule inhibitors in SCLC, but not NBL cell lines, whereas higher NE scores were associated with resistance to BRD inhibitors in NBL, but not SCLC lines (**Figure 6**). Notably, although we identified differential therapeutic sensitivity within SCLC and NBL panels relative to their NE lineage, this does not tell us about the dynamic ranges of compound sensitivity in SCLC and NBL. In some cases, the dynamic range of compound sensitivity remains different between SCLC and NBL. For example, SCLC cell lines are the most resistant to MEK inhibitors, whereas NBL cell lines exhibit intermediate sensitivity over a broader range (**Figures S6a**-**c**). In other cases, we observed a similar overall sensitivity of SCLC and NBL cell lines. For example, both SCLC and NBL cell lines were more resistant to the HSP90 inhibitor 17-AAG, but more sensitive to the BCL inhibitor ABT-199, compared to the other cancer lineages (**Figures S6d-e**). In summary, our results revealed that the relative differential drug sensitivities associated with NE-to-non-NE transdifferentiation in SCLC and NBL were similar.

**Figure 6.**
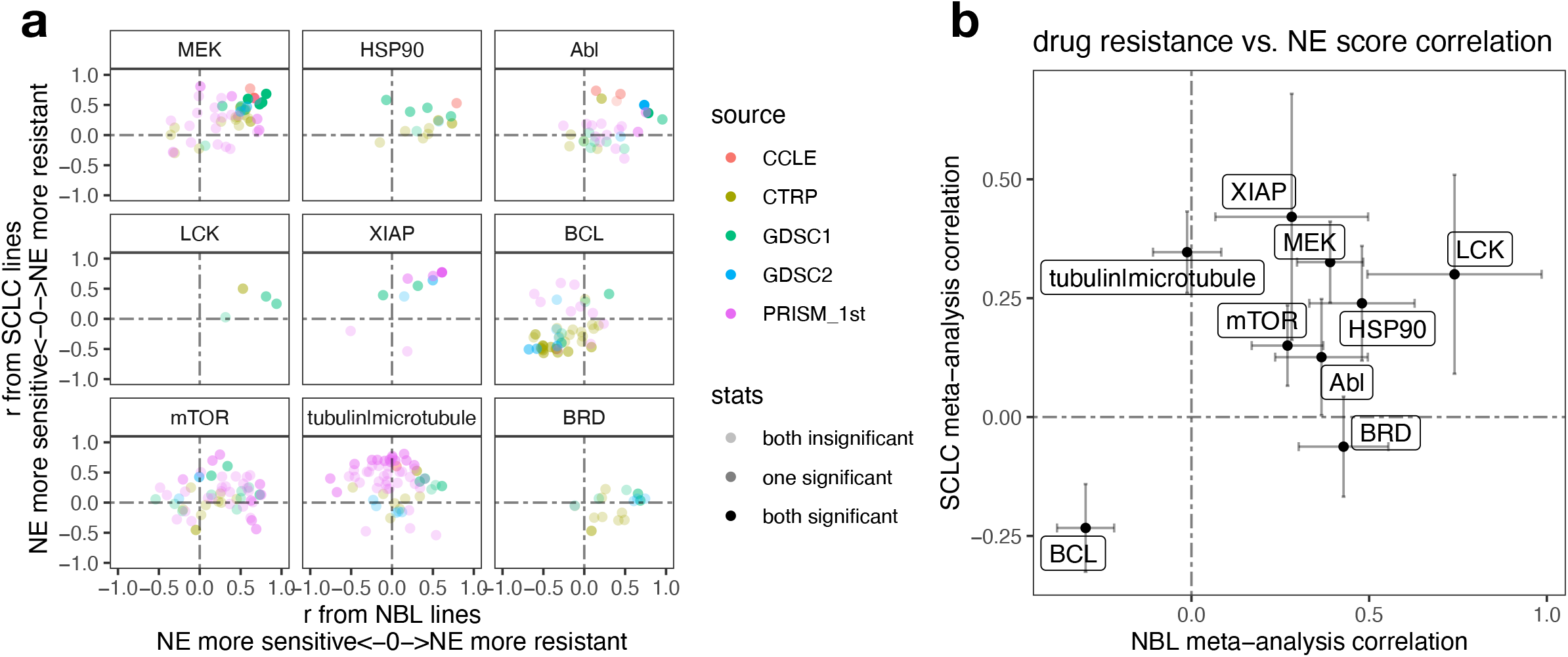
Similar and distinct NE score-associated therapeutic sensitivity in SCLC and NBL cell lines. **a**. Correlation between NE scores and therapeutic sensitivity for drugs with selected targets. Therapeutic sensitivity data was previously harmonized such that a higher value represents more resistance in each study. For each of the nine selected targets, all compounds with the same target were identified from multiple studies. Pearson correlation coefficient r from correlating compound data with NE scores were calculated for NBL lines (x-axis values) and SCLC lines (y-axis values) respectively and visualized as a scatter plot, with colors annotating the source of data, and transparency annotating the statistical significance. **b**. Meta-analysis-summarized correlation between drug therapeutic sensitivity and NE scores in NBL (x-axis) and SCLC (y-axis) cell lines. Note that high NE scores are associated with resistance to inhibitors of LCK, MEK, XIAP, mTOR, HSP90, and Abl, and sensitivity to BCL inhibitors. NE scores are associated with resistance to BRD inhibitors in NBL but not SCLC whereas microtubule inhibitors resistance correlates with high NE scores in SCLC but not NBL cell lines.

### Identification and comparison of SCLC and NBL-specific gene dependencies

We observed very poor overall concordance between the RNAi and CRISPR dependency data for their association with NE scores in SCLC and NBL (**Figure S4**). We rationalized that this is because most genes were not selectively essential in the relatively small panel of SCLC or NBL cell lines assessed. Hence, we adopted a set of criteria for selecting cancer-specific gene dependencies. We looked for genes with RNAi vs. CRISPR gene effect scores positively correlated, as an indication of high reproducibility from independent dependency screening experiments, as well as negative correlations between RNAi or CRISPR gene effect scores and RNA-seq expression data on the premise that genes of selective functional importance are more highly expressed in the cells that depend on them, such that these cells also have more negative gene effect scores that indicate higher dependence. These measures from the SCLC and NBL panels were assembled to prioritize the SCLC-specific vulnerabilities (**Table S13**). Indeed, when we examined these correlations, the known SCLC subtype drivers and the most common NBL driver genes all met this set of criteria (**Figure 7a-b**). We further closely examined genes with high RNAi vs. CRISPR correlation, and high anti-correlations between RNA expression and the gene effect scores as selected vulnerabilities (**Figure 7c**). Among the SCLC-selected vulnerabilities, along with ASCL1, we found several other NE lineage transcription factors (*SOX11, FOXA2, NKX2-1*) were more selectively essential for high-NE-score cell lines, whereas several genes involved in cell adhesion and motility (*VCL, PXN, ACTR3*, and *RAC1*) were found to be more selectively essential for low-NE-score cell lines; we also found genes frequently amplified in SCLC (*IRS2, CCNE1*, and *NFIB*) (Kim et al., 2019) although these genes do not have gene effect scores significantly correlated with NE scores. Interestingly, among these SCLC-selected vulnerabilities, we also identified genes that are well characterized for their roles in NBL, such as the ciliary neurotrophic factor *CNTF* (Heymanns and Unsicker, 1987) and S-phase kinase-associated protein 2 *(SKP2* (Muth et al., 2010). Among the very few vulnerabilities selected from both SCLC and NBL, we identified *BCL2*, a well-characterized gene in both cancer types. Consistent with our observation in the therapeutic sensitivity analysis, high-NE-score cell lines from both SCLC and NBL were more sensitive to *BCL2* depletion (**Figures 7d-e**). As only nine NBL cell lines were included in the RNAi dependency screen, the reliability of our NBL-selected vulnerabilities might have been undermined by the underpowered input datasets. Nevertheless, we were able to identify a few genes known to be important for NBL, such as GATA Binding protein 3 *GATA3* (Hoene et al., 2009), complement decay-accelerating factor *CD55* (Weng et al., 2022), Forkhead Box R2 *FOXR2* (Schmitt-Hoffner et al., 2021), and breast cancer anti-estrogen resistance protein 1 *BCAR1* (/p130Cas) (Fagerstrom et al., 1998). Among these, selective essentiality for *GATA3* was only observed for high-NE-score NBL cell lines but not SCLC cell lines (**Figure 7f-g**), whereas *BCAR1* appears to be a shared vulnerability for low-NE-score cell lines in both NBL and SCLC (**Figures 7h-i**). Overall, we observed unique and shared gene dependencies between SCLC and NBL cell lines, some of which also exhibited NE/non-NE lineage-specific selectivity.

**Figure 7.**
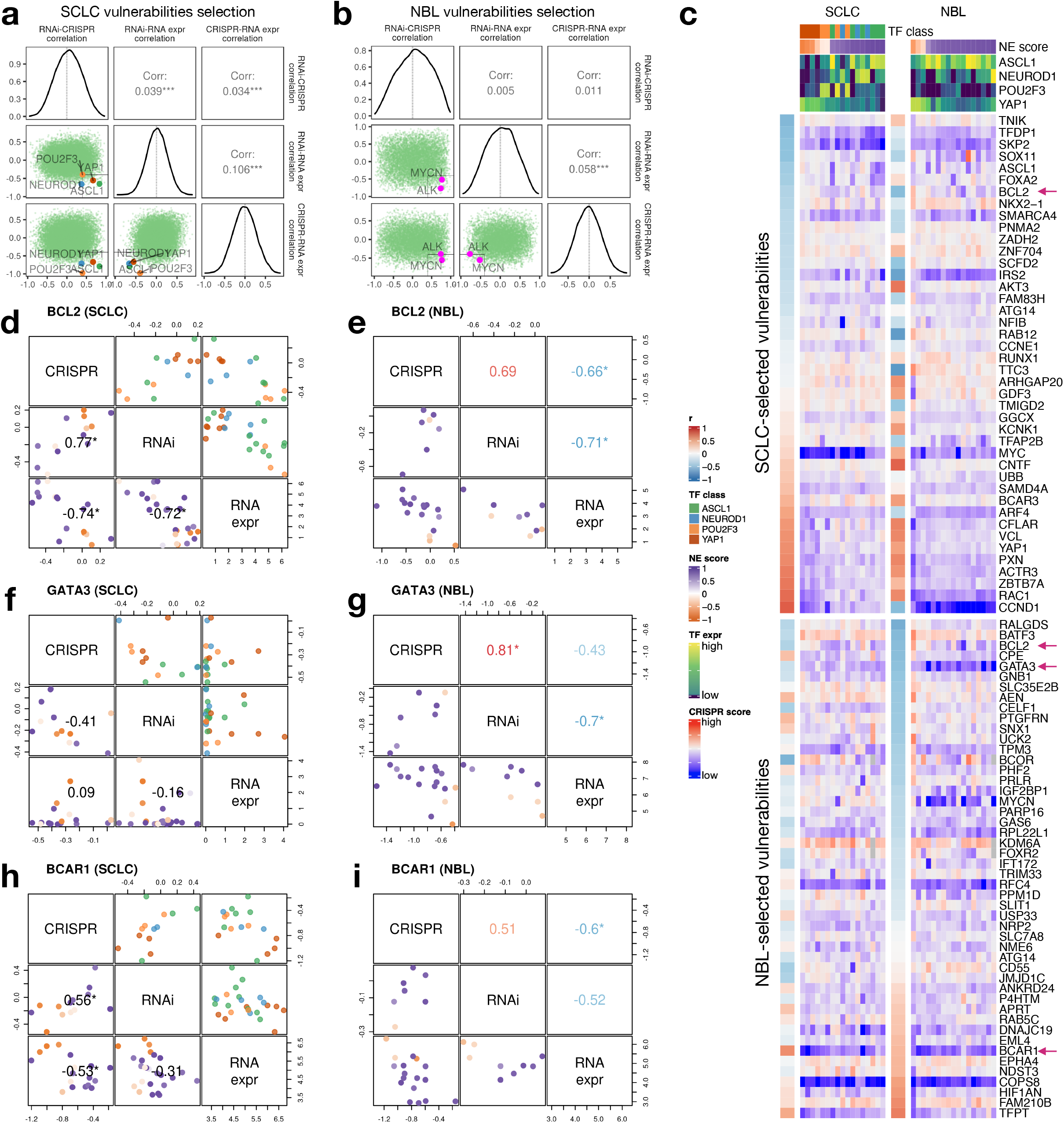
Similar and distinct NE score-associated gene dependencies in SCLC and NBL cell lines. **a-b**. Selection of SCLC (**a**) and NBL (**b**) vulnerabilities based on the consistency (positive correlation) between CRISPR and RNAi data, and anti-correlation between dependency data and gene expression data. Pearson correlation coefficients from RNAi-CRISPR (left), RNAi-RNA expr (middle), and CRISPR-RNA expr (right) correlations were computed for all genes. The distributions of these coefficients are plotted as diagonal panels; pairwise correlations among these three sets of correlation coefficients were visualized as scatter plots in the lower triangular panels and the Pearson correlation coefficients are printed in the upper triangular panels. The four SCLC subtype driver TFs and the NBL oncogenic driver MYCN all have high consistency between CRISPR and RNAi data and high anti-correlation between dependency data and gene expression data. Areas with r > 0.4 from RNAi - CRISPR correlation, and r < −0.4 from RNAi - RNA expr and CRISPR - RNAi correlation were demarcated by light gray squares. **c**. Correlation between NE scores and effect scores of selected dependencies in SCLC and NBL. The upper part of the heatmap displays selected vulnerabilities for SCLC and was ordered by correlations between NE scores and the effect scores in SCLC cell lines; likewise, the lower part of the heatmap displays selected vulnerabilities for NBL. Genes with magenta arrows are showcased in **d**-**i**. Cell lines are ordered by their NE scores and annotated with NE score and SCLC driver TF expression. **d**-**i**, Comparison of selected gene dependencies in SCLC and NBL. In each plot, variable names are shown in the diagonal boxes, and scatter plots display relationships between each pairwise combination of variables. Lower triangular plots are colored by NE scores whereas upper triangular plots for SCLC figures are colored by TF classes. Pearson correlation coefficients are provided in lower triangular boxes for SCLC and upper triangular boxes for NBL. Refer to legends in **c** for color annotations.

## Discussion

Different cancers of the NE lineage have historically been investigated as separate entities, owing to their distinct clinical presentations. The common Notch-mediated NE lineage plasticity and the adjacency of SCLC and NBL in cancer cell line clustering by multi-omics datasets (**Figure 1**) prompted us to perform a systematic comparison of these two cancer types. In this paper, we identified numerous common molecular associations with NE states in both cancer types. Most of the proteomic and metabolic features observed to associate with NE states in SCLC could be validated in NBL (**Figure 5**). NE score-associated transcriptomes are also highly similar between SCLC and NBL. We previously reported cell-autonomous immune gene repression in SCLC and pulmonary neuroendocrine cells in the NE state and transdifferentiation into the non-NE lineage releases the repression of immune genes (Cai et al., 2021a). Recently, similar observations have been reported for NBL (Sengupta et al., 2021). Besides immune genes, many other genes are differentially expressed by NE status in both SCLC and NBL. Although our omics analyses in this study do not include large-scale transcriptomics comparison, this topic is explored in greater depth in a companion manuscript (Cai et al., 2022).

Our comparison of SCLC and NBL therapeutic vulnerabilities revealed similar NE score associations for several classes of compounds with shared MOA. Interestingly, many of these have also been reported in the literature by individual studies in SCLC or NBL. For example, MEK inhibitors for NBL (Tanaka et al., 2016), HSP90 inhibitors for SCLC (Workman and Powers, 2007) and NBL (Regan et al., 2011), BCL2 inhibitors for SCLC (Fennell, 2003) and NBL (Lamers et al., 2012), etc. We also identified cancer-unique vulnerabilities that have been previously reported, for example, BRD inhibitors and GATA3 essentiality for NBL (Peng et al., 2015; Puissant et al., 2013). Importantly, our findings that these functional liabilities strongly correlate with NE scores suggest NE plasticity may serve as a venue for therapy resistance in both SCLC and NBL for such drugs, as long-term monotherapy that targets cells of one lineage might shift the population towards the other lineage. Coupled with our observation that high- and low-NE-score cells can co-exist within the same SCLC or NBL cell line (**Figure 2**), it would be interesting to compare the efficacy of monotherapy to combinatorial therapies that target both NE and non-NE states using cell lines or cell line-derived xenografts as preclinical models.

Compared to the omics analyses, relatively fewer similarities were observed at the functional liability levels. Several reasons might explain this discrepancy. First, functional data is much noisier than -omics data (Cai et al., 2021b). Second, fewer SCLC and NBL cell lines were included in the functional screening datasets and this compromised the statistical power for target discovery (**Figure S4**). As our concordance-based approach requires examining data from common cell lines between two datasets, this further reduces the available sample size for analysis. Lastly, the unique vulnerabilities for SCLC and NBL may stem from cancer drivers that act orthogonally to NE status. One such example is *NFIB*, which has been characterized as a metastatic promoter in SCLC (Semenova et al., 2016).

In summary, by comparing SCLC and NBL side by side, we created a comprehensive reference for molecular features and vulnerabilities commonly associated with NE-to-non-NE lineage transitions. Our analyses also revealed SCLC- and NBL-unique features that need to be investigated in cancer-specific contexts. These results may serve as a reference for designing combinatorial therapies targeting lineage plasticity in SCLC, NBL, and other NE cancers.

## Material and Methods

### Input datasets

#### Cell line datasets

Copy number, RNA-seq, miRNA, histone PTM, metabolomics, and RPPA data were downloaded from DepMap. Compound sensitivity data for “CCLE” (Barretina et al., 2012), “CTRP”(Basu et al., 2013), “GDSC1” and “GDSC2” (Iorio et al., 2016), “PRISM_1st” and “PRISM_2nd” (Corsello et al., 2020), and gene dependency data for demeter (RNAi) (McFarland et al., 2018) and achilles (2020) (CRISPR) were downloaded from DepMap and processed as previously described (Cai et al., 2021b). The cell line names and compound names were unified, and the datasets were processed to ensure that the lower value in each dataset always corresponded to a higher sensitivity. The processed data, lists of consistent compounds, and dependencies were downloaded from https://lccl.shinyapps.io/FDCE/. The scRNA-seq data for cell lines were downloaded from the Gene Expression Omnibus (GEO) repository GSE157220 (Kinker et al., 2020).

#### Additional SCLC datasets

The following SCLC transcriptomic datasets “UTSW SCLC cell line,” “Drapkin_2018” (PDX) (Drapkin et al., 2018), tumor datasets “Rudin_2012” (Rudin et al., 2012), “George_2015” (George et al., 2015), “Jiang_2016” (Jiang et al., 2016), and “Cai_2021” (Cai et al., 2021a) were processed as previously described (Cai et al., 2021a). The processed data are available in our previous publication (Cai et al., 2021a). SCLC scRNA-seq data were downloaded from the HTAN portal (Chan et al., 2020).

#### Additional NBL datasets

In addition to the CCLE RNA-seq data, additional NBL cell line transcriptomic and associated sample phenotype data were downloaded from GEO using R package GEOquery (Davis and Meltzer, 2007) with the following accession numbers: GSE28019, GSE89413 (Harenza et al., 2017), and GSE90683 (Boeva et al., 2017). For NBL patient tumor datasets, we included two partially overlapped NBL datasets from Therapeutically Applicable Research to Generate Effective Treatments (TARGET) (https://ocg.cancer.gov/programs/target) initiative, phs000467 (Pugh et al., 2013). “TARGET_microarray” was downloaded from the TARGET Data Matrix, whereas “TARGET_RNA-seq” was downloaded from the UCSC Toil RNAseq Recompute Compendium (Vivian et al., 2017). Additional NBL tumor datasets were downloaded from GEO with the following accession numbers: GSE120572 (Ackermann et al., 2018), GSE3446(Asgharzadeh et al., 2006), GSE19274(Cole et al., 2011), GSE73517(Henrich et al., 2016), GSE85047(Rajbhandari et al., 2018), GSE62564(Wang et al., 2014), GSE16476(Molenaar et al., 2012), and GSE3960(Wang et al., 2006).

### Clustering of cell lines by multiomics, drug sensitivity, and dependency data

For the clustering of cell lines based on RNA-seq data, we first conducted a principal component analysis for genes with a standard deviation larger than 0.4. The top ten principal components accounted for 41% of the total variance and were used for hierarchical clustering. For the clustering of RPPA and metabolomics data, we used all available features and did not filter the input features or perform principal component analysis. For clustering dependency and drug data, we used our previously generated consistency measures based on meta-analysis-summarized inter-study pairwise Pearson correlations (Cai et al., 2021a). For dependency data, we selected consistent features with r > 0.4, and for drug data, we selected consistent features with multiple comparison-adjusted p-values < 0.05. All hierarchical clustering was performed using Ward’s minimum variance method.

### NE score computation

The original SCLC NE signature based on microarray gene expression data was described by Zhang et al. (Zhang et al., 2018). Here, we used the updated signature generated from RNA-seq expression(Cai et al., 2021a). A quantitative NE score can be generated from an NE signature using the formula NE score□=□(correl NE – correl non-NE)/2, where correl NE (or non-NE) is the Pearson correlation between the expression of the 50 genes in the test sample and expression/weight of these genes in the NE (or non-NE) cell line group. This score ranges from −1 to +1, where a positive score predicts NE and a negative score predicts non-NE cell types. The higher the score in absolute value, the better the prediction.

## Acknowledgment

L.C. received support from UTSW ACS-IRG (IRG-21-142-16) and a Lung Cancer SPORE Career Enhancement Program award from P50CA70907. This study was supported by funding from the National Institutes of Health [R35CA22044901, P30CA142543, P50CA70907, and R35GM136375], and the Cancer Prevention and Research Institute of Texas [RP190107 and RP180805]. R.J.D. received funding from Howard Hughes Medical Institute. This article is subject to HHMI’s Open Access to Publications policy. HHMI lab heads have previously granted a nonexclusive CC BY 4.0, license to the public, and a sublicensable license to HHMI in their research articles. Pursuant to those licenses, the author-accepted manuscript of this article can be made freely available under a CC BY4.0 license immediately upon publication.

