## supplementary figures 1-6 for "A comparative study of neuroendocrine heterogeneity in SCLC and NBL"

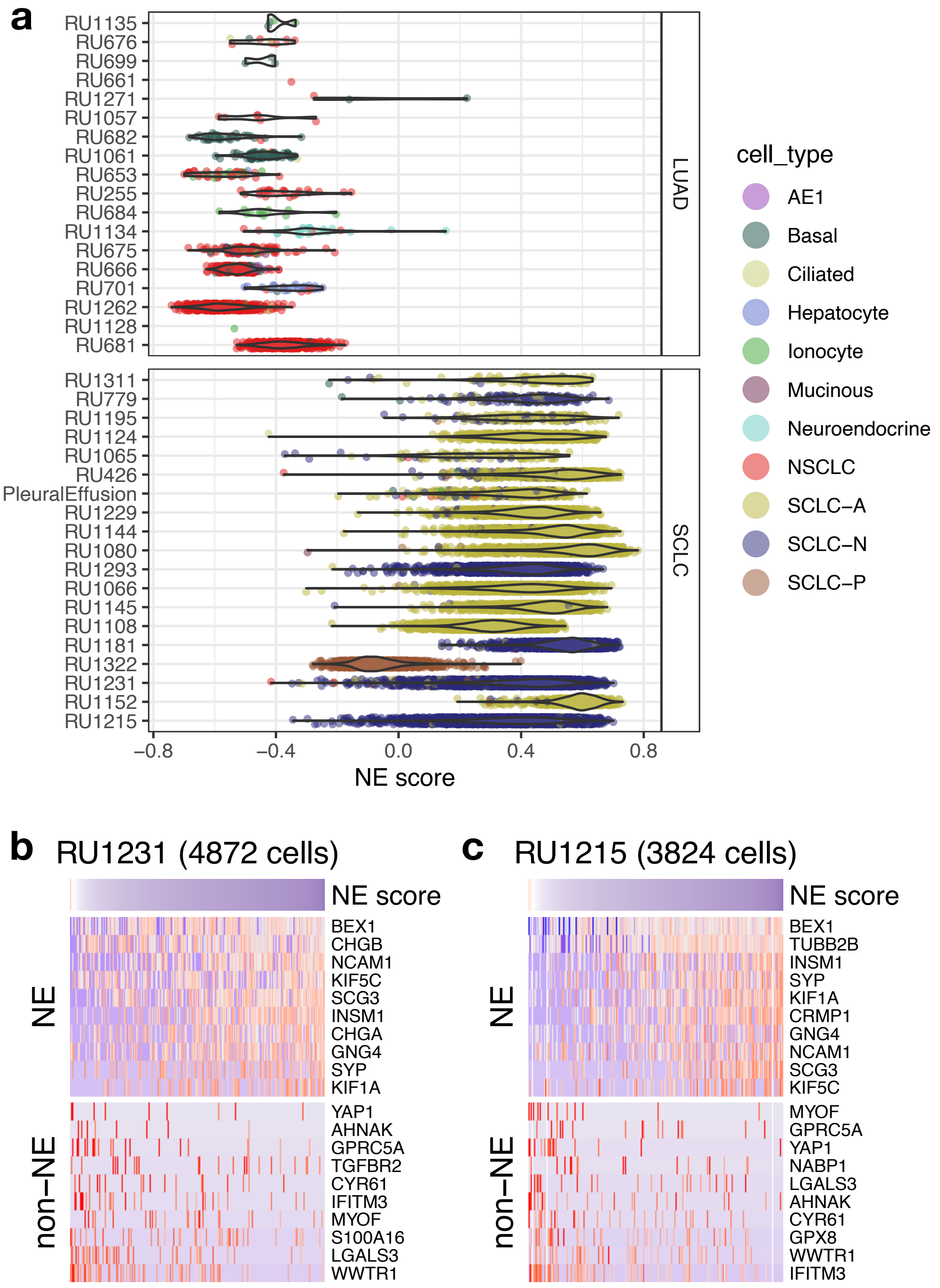


**Figure S1. Intratumoral NE heterogeneity in SCLC patient tumors**

**a**. Distribution of NE scores for lung cancer patient tumors from HTAN datasets. Sample names are labeled on the y-axis. Each dot represents a cell within the sample and is colored by the cell type annotated by the original study, violin plot is overlaid to visualize the overall distribution. Note that two SCLC-N tumors (RU1231 and RU1215) have broad NE score distributions. **b-c**. High- and low- NE score tumor cells are found to coexist within the same SCLC tumor. Single cells are in columns. Due to the high dropout rate of scRNA-seq data, only the top abundantly expressed genes are visualized.

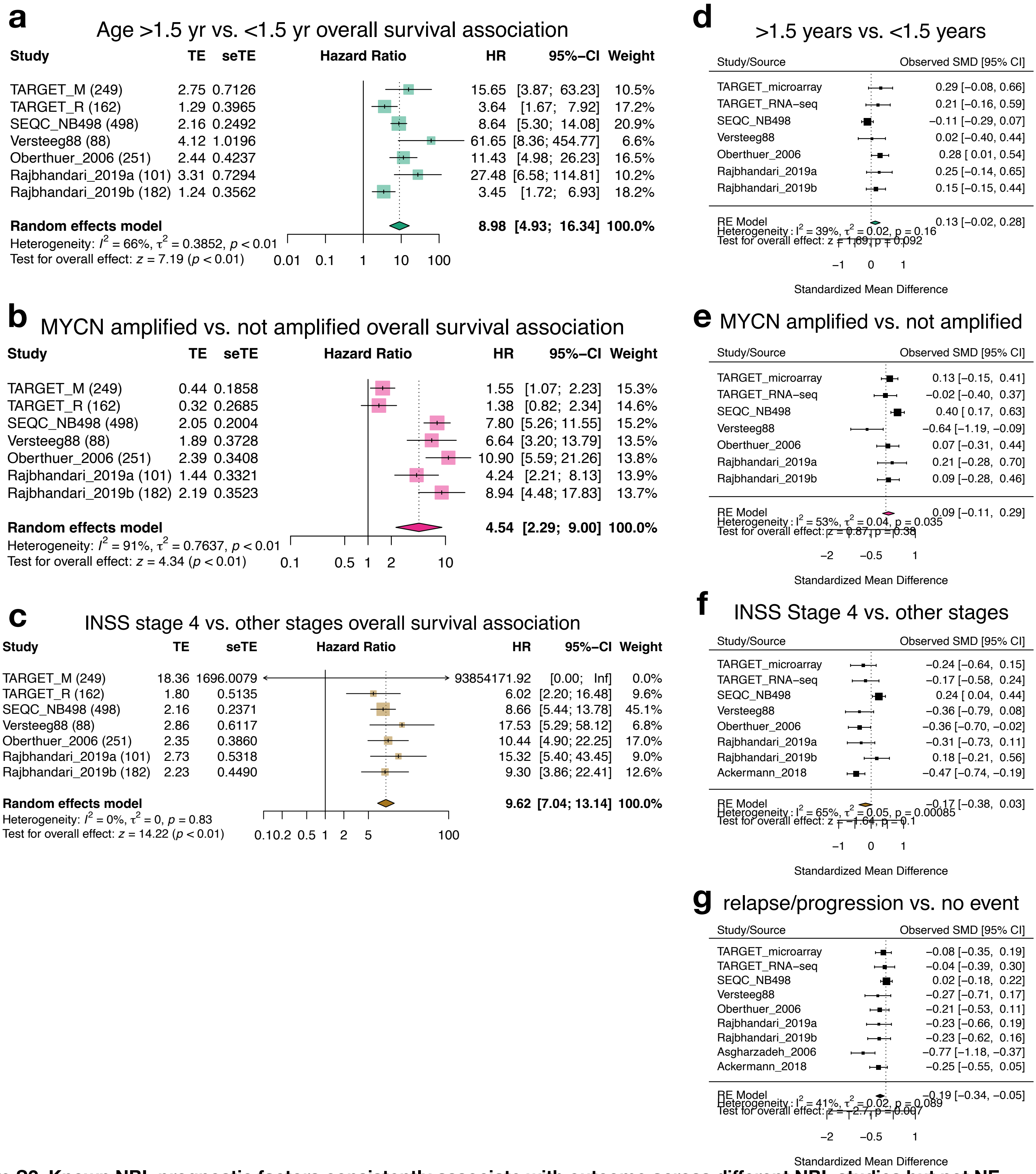


**Figure S2. Known NBL prognostic factors consistently associate with outcome across different NBL studies but not NE scores**

**a-c.** Age at diagnosis (**a**), MYCN amplification (**b**), and INSS stage 4 (**c**) are significantly associated with worse outcome. **d-g**. Meta-analyses of NE scores standardized mean difference between groups stratified by different clinical features. Age at diagnosis (**d**), MYCN amplification (**e**) and INSS stage 4 (**f**) are not significantly associated with NE scores. Lower NE scores are associated with relapse (**g**). Note that although one out of the nine studies showed a significant result, the overall result from the meta-analysis is statistically significant.

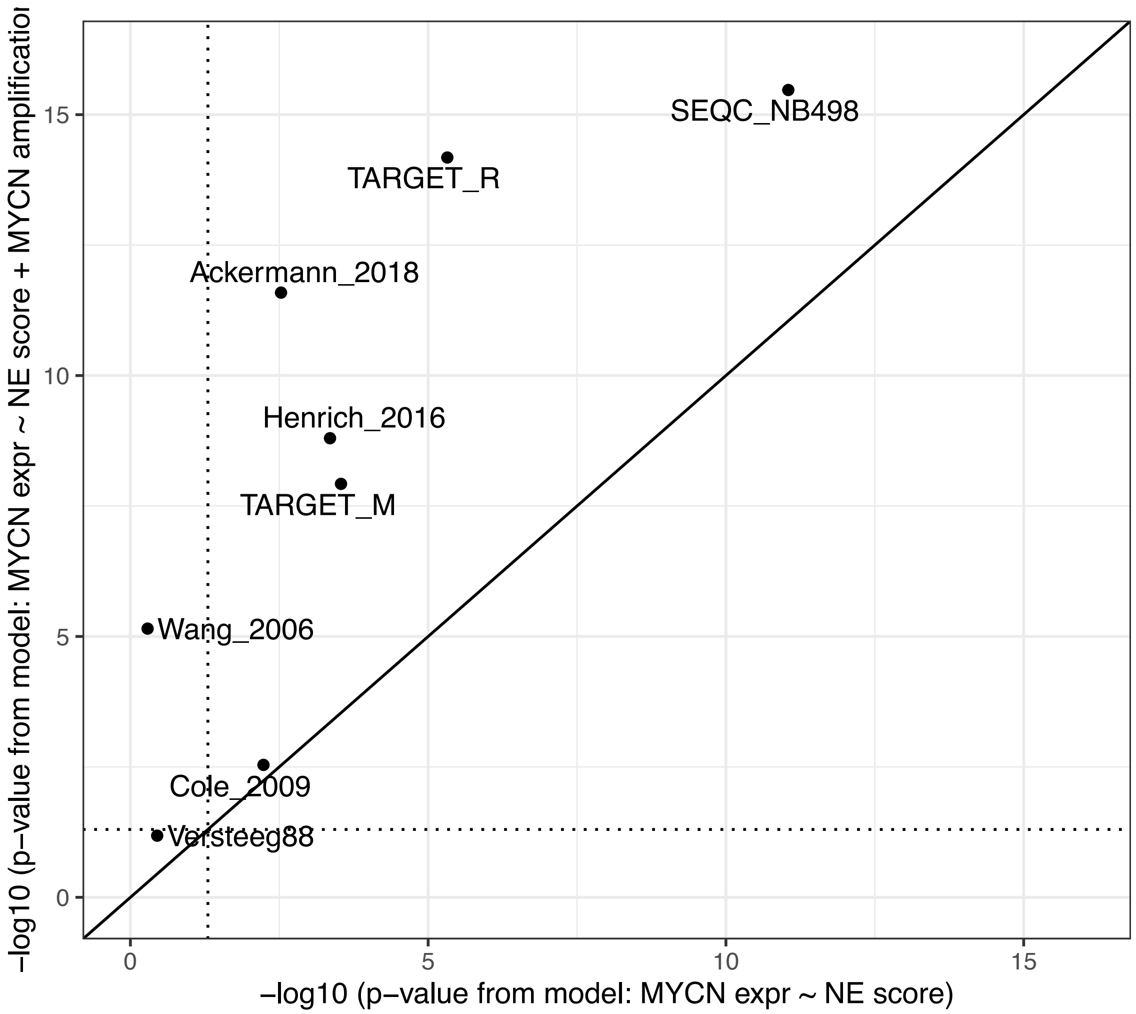


**Figure S3. Controlling for MYCN amplification status increases the statistical significance of NE score vs. MYCN expression association.**

X-axis values are p-values from a univariate linear model using NE score to predict MYCN expression, y-axis values are p-values from a multivariate linear model that included both NE score and MYCN amplification status as the predictor variables and MYCN expression as the response variable. For both models, p-values for the NE score term were extracted for comparison. Results from eight studies were included and study names were labeled on the plot. Dotted line indicates p-value = 0.05.

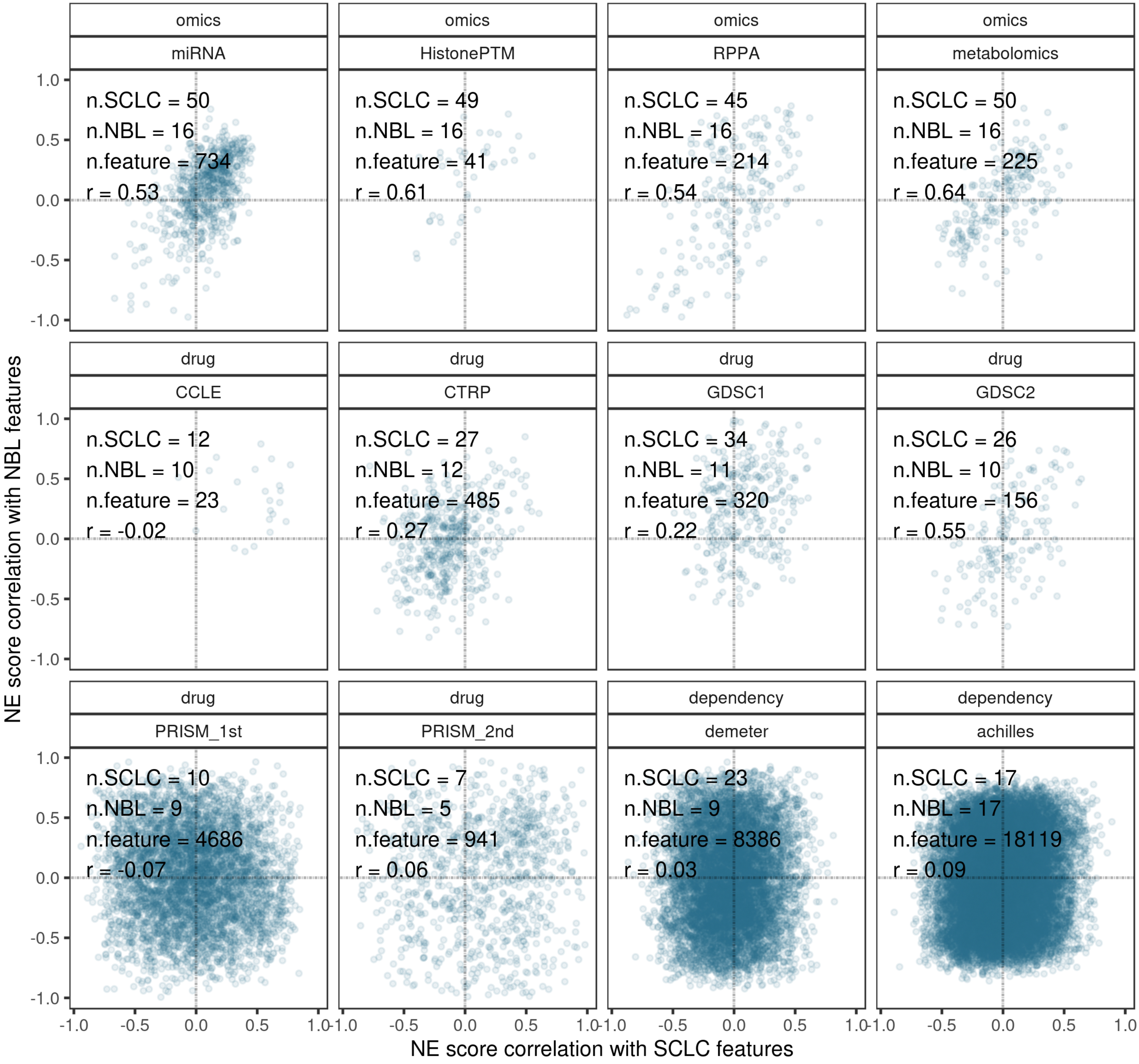


**Figure S4. SCLC vs. NBL concordance of NE score association with omics, drug sensitivity, and dependency data**

For each dataset, we correlated each feature with NE scores in SCLC cell lines to obtain a set of correlation coefficients (x-axis values); we also generated such values for NBL cell lines (y-axis values). Correlation from correlating SCLC results and NBL results was further conducted to summarize an overall concordance metric “r” for each dataset. Parameters printed at the upper left quadrant of each plot: n.SCLC, the median number of SCLC cell lines in correlation; n.NBL, the median number of NBL cell lines in correlation; n.feature, the number of features in the dataset; r, Pearson correlation coefficient from correlating x-axis and y-axis values, representing concordance between SCLC and NBL NE score correlations.

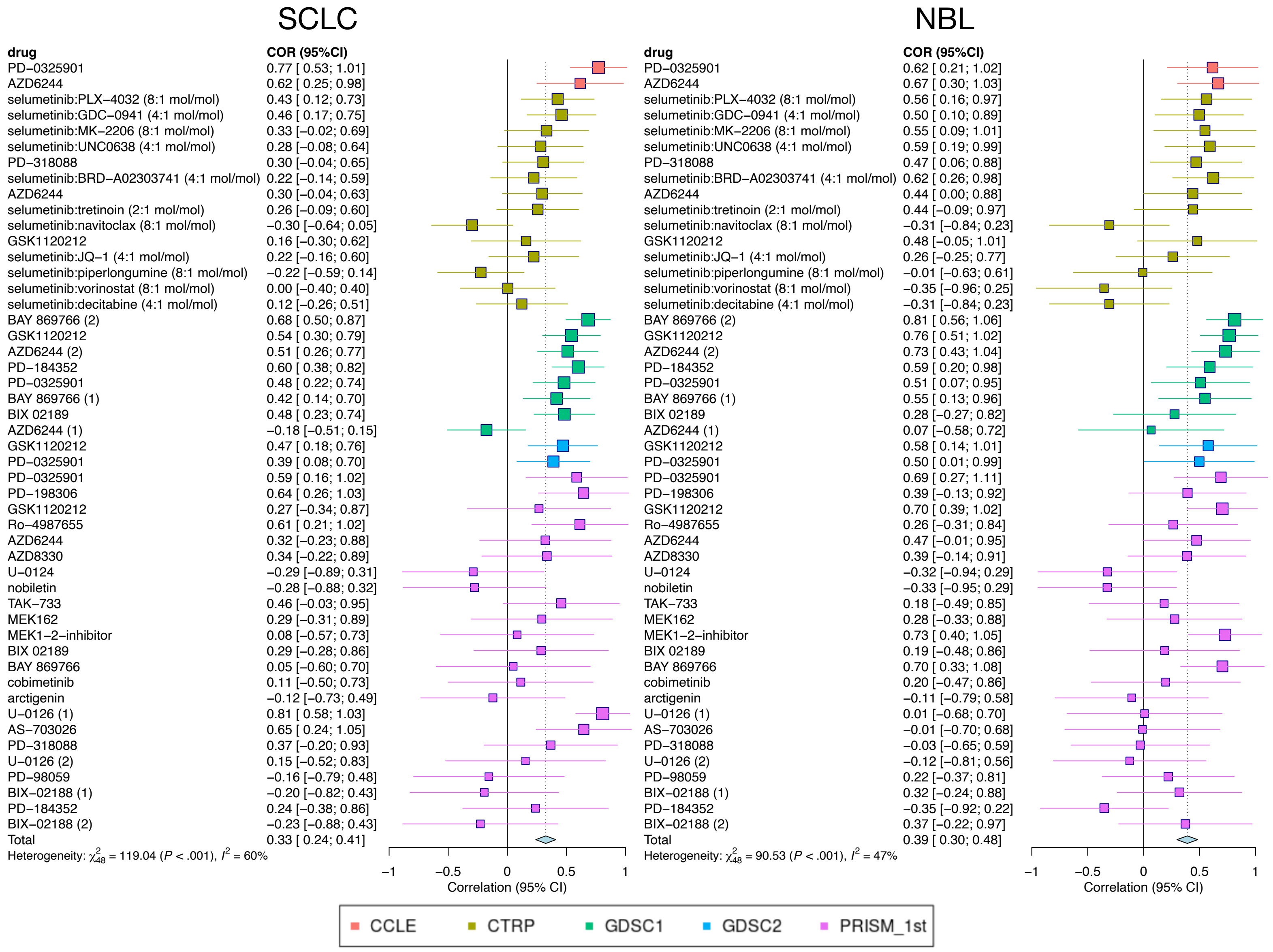


**Figure S5. A meta-analysis example that summarizes the association between MEK inhibitor resistance and NE scores**

Compounds that target MEK were identified from five different datasets. Correlation between therapeutic sensitivity of different MEK inhibitors and NE scores in SCLC and NBL cell lines were computed. These forest plots visualize the meta-analyses of these correlations in SCLC (left) and NBL (right). Sources of data were annotated in different colors. Datasets have been harmonized such that a positive correlation indicates cell lines with higher NE scores are more resistant.

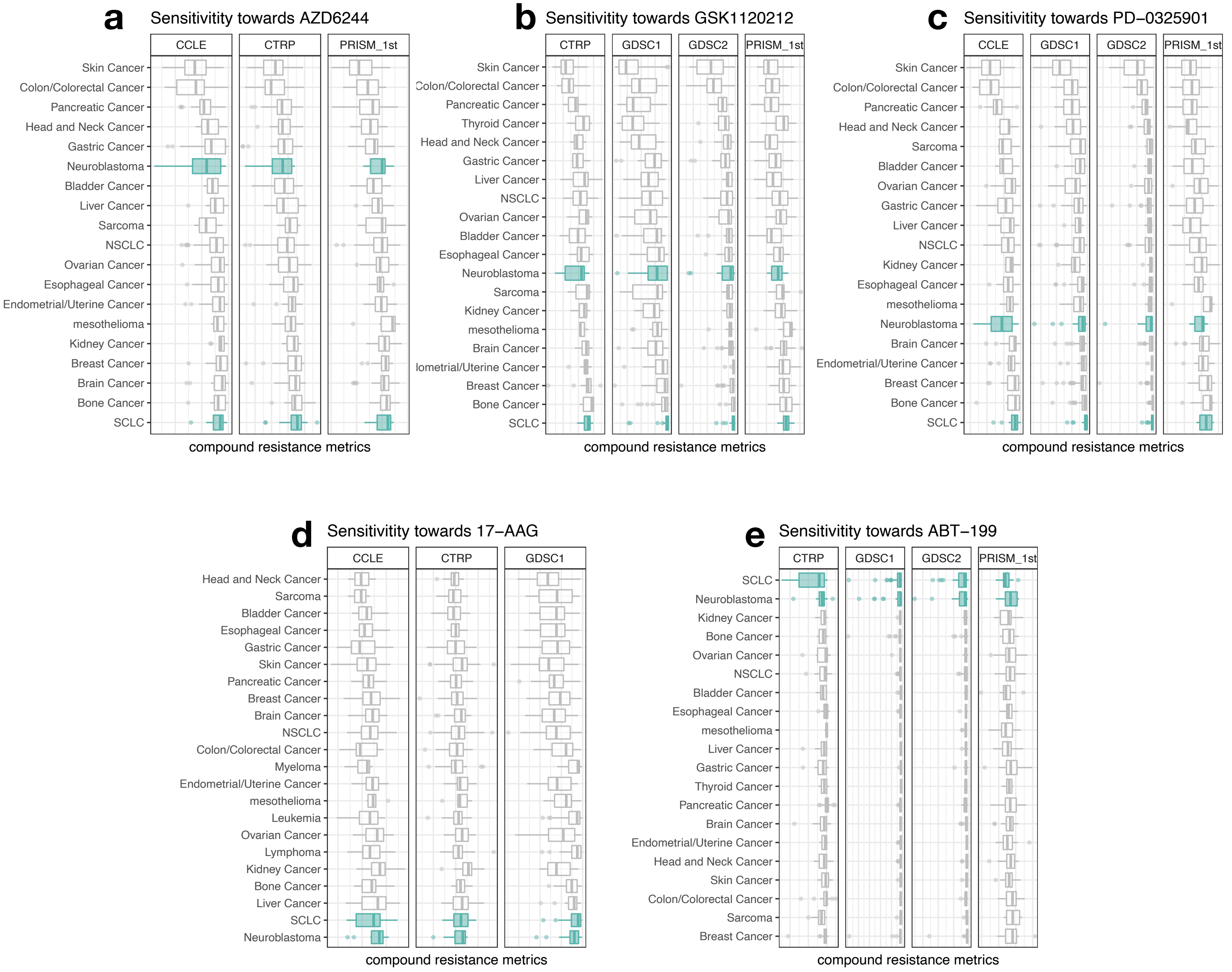


**Figure S6. Examples of different and similar compound sensitivity dynamic ranges in SCLC and NBL**

**a-e**, Distribution of compound sensitivity data by cancer lineage for selected MEK inhibitors (**a-c**), HSP90 inhibitor (**d**), and BCL inhibitor (**e**). Compound resistance metrics were z-transformed within each study. For each compound, the cancer lineages were ordered by median compound resistance values from all studies. Note that skin cancer (primarily melanoma) cell lines are most sensitive to MEK inhibitors whereas SCLC lines are most resistant to MEK inhibitors. NBL cell lines exhibit a broad distribution of sensitivity and have intermediate sensitivity to MEK inhibitors compared to the other cancer types (**a-c**). SCLC and NBL exhibited more similar compound sensitivity ranges for 17-AAG (**d**) and ABT-199 (**e**).
